# Identification of the combining abilityof grain yield and its components in hybrid barley populations based on topcross analysis

**DOI:** 10.1101/2021.04.13.439609

**Authors:** Akhmedova Gulmira, Tokhetova Laura, Umirzakov Serikbai, Demesinova Ainur, Tautenov Ibadulla, Bekzhanov Serik, Kali Omarov, Shayanbekova Bakhytzhan

## Abstract

The topcross method for assessing combining abilityis more economical and less laborious compared to diallel analysis, and also allows the breeder to obtain quite valuable information about the inbred material. In this research, the determination of the general (GCA) and specific (SCA) combining ability of barley (Hordeum vulgare L.) hybridsin two regions of Kazakhstan contrasting in soil and climatic conditions,the role of additive and non-additive genes in the determination of thetraits understudyhas beenrevealed. It is concluded thatthe predominance of additive gene interactions in the control of the traits understudyin theconditions of the Aral Sea region indicates the possibility of effective selection already in the F_2_ generation, and in the piedmontzone of the Almaty region, due to the high determination of these traits by dominant genes, it is necessary to differentiate the populations of hybrids, starting from the first generation and further selection shall be carried out in several cycles until the achievement of homozygosity of loci carrying dominant genes. Consequently, the genetic contribution of the additive and non-additive effects of genes to the determination and inheritance of the studied traits significantly depends on the conditions of growing the genotypes of spring barley. Of greatest practical interest are the varieties Rihane, WI2291/Roho/WI2269 from the International Center ICARDA and the variety-tester Odessa 100 (Odessa Selection and Genetic Institute, Ukraine) with high GCA and SCA effects, little dependent on growing conditions, which are recommended to be used as reliable donors in breeding programs.

## 1. Introduction

The Kyzylorda region of the Republic of Kazakhstan is geographically located in extremely unfavorable conditions for the implementation of crop production, where there is a decrease in the water resources of the transboundary Syrdarya River, which creates certain threats in ensuring guaranteed water supply to irrigated lands, causing intense desertification, salinization and soil blowing, which are the main obstacle to sustainable economic growth and social development of the region. In this regard, within the framework of the “green economy” and the diversification of crop production in the Kyzylorda region, the areas under low-water consumption agricultural crops are expanding, including the fodder-grain crop – barley, which is one of the leading crops in the world, due to its adaptive capabilities, high yield and versatile use [1, 2]. However, the production of own fodder fails to meet the needs of the cattle breeding of the Kyzylorda region, it has to be purchased additionally, one of the reasons for which is the low yield of agricultural crops on saline soils. Therefore, one of the main tasks of selection work on barley in this region is to search for high yielding adaptive genotypes that are stable by year, and to determine their donor properties to attract them to hybridization programs.

The effectiveness of selection largely depends on the genetic analysis of the hereditary qualities of the source material. Genetic counseling, preservation of the genetic health of the population, protection of the existing gene pool are also ultimately based on genetic analysis. Therefore, further development and improvement of methodsof genetic analysis continue to be the most important task of modern genetics [3, 4]. It is very important to identify from a variety of source material forms that not only combine a valuable complex oftraits, but also have the ability to breed true, to select combinations of crossing in which the desired transgressions can be obtained. One of the methods of analysis that allows to evaluate the genetic properties of varieties is to identify the combining ability, which makes it possible to the researcher to anticipate the results of future crosses and focus on promising material, while avoiding unnecessary time and money outlays on re-obtaining and testing hybrids of no practical value. To determine the general (GCA) and specific combining ability (SCA) during cross-pollination, both the diallel scheme [5-8] and the system of topcrossbreeding [9-11] are used. The topcross method for assessing combining ability is more economical and less laboriouscompared todiallel analysis, and also allows the breeder to obtain quite valuable information about the inbred material.

Literature review has shown that different genes are involved in the determination of traits, sometimes they have a contradictory character in the limit of one agricultural crop, which, apparently, depends on the genotype of the parental lines involved in hybridization and, in the case of polygenic inheritance, can be also determined by the growth conditions of plants [12-14].

## 2. Materials and Methods

The research was carried out simultaneously in two zones of Kazakhstan, contrasting in soil and climatic conditions: - rice systems of the Kyzylorda region; - piedmont zone of the Almaty region.

The climate of the Kyzylorda region is sharply continental with hot dry summers and cold winters with unstable snow cover. The average annual air temperature is 9.8^0^C. The climate of the region is very dry. The average annual precipitation is 129 mm. In some dry years, precipitation can be 40-70 mm. The soil of the experimental field is meadow-boggy, typical for rice crop rotations in the region. It has a low humus content of 0.9% and a high value of dissolved solids – 0.88%. Salinizationis average,and chloride-sulfate.

The climatic conditions of the piedmont zone of the Almaty region are characterized by cold winters, hot and dry summers, warm and dry autumns. Average air temperature 7.6^0^C. The average annual precipitation is 414 mm. The soil cover is represented by light chestnut (nonsaline) soils. The humus content reaches 3%.

The topcross method is widely used to assess the combining ability of lines. The essence of the method lies in the fact that all lines being studied are crossed with a common tester. Lines, hybrids or varieties can be used as a tester, and, as a rule, there should be at least two testers. Testers can be used as both female and male parents. The more testers are used in crossing, and the more genetically diverse they are, the more accurate the assessment of the general and specific combining ability will be. The stress factors of the environment (salinity, drought, and dryhot winds), characteristic of the soil and climatic conditions of the Kyzylorda region, greatly reduce the setting of grain during hybridization and averages 3-5%.Therefore, it is very difficult to obtain the entire set of planned hybrid populations in diallel crossings. Many hybrid combinations “fall out” and it is not possible to carry out a genetic analysis. In this regard, in our research we widely use the topcross method, based on which we have created not only valuable genetic material, but also salt-tolerant high-yielding fodder barley varieties that are in demand withagricultural commodity producers in the country. In this research, using the topcross method for F_1_ hybrids, the general (GCA) and specific (SCA) combining ability of barleyvarieties were studied: WI2291*2/WI2269; WI2291/Roho/WI2269; Rihane, Harmal (ICARDA) used as female parents. Variety-testerswere used as male parents: Donetskiy8, Odessa 100 (Ukraine), andSaule (Kazakhstan). The main criteria for the selection of female parents were early maturity anddwarfness; of male parents– high grain content, grain size and tallness.

Seeds were sown on 1-meter rows with 15 cm row spacing in three repetitions. F_1_ plants were harvested by hand together with roots. 50 plants were analyzed according to individual quantitative traits: productivity (grain weight) of a plant, its structural elements (productive tilling capacity, number of grains perspike, thousand grain weight), as well as plant height and spike length. The reliability of impact of factors on the variability of combining ability indicators under the influence of environmental factors was assessed by the method of Tarutina A.I. and KhotylevaL.V. [15]. The analysis of combining ability in topcrossbreeding was carried out by the method of V.K. Savchenko [10].

## 3. ResultsandDiscussion

Analysis of variance showed the presence of a significant variation in the studied traits depending on the testingsites with the exception of the generalcombining abilityof the grainweight per plant of male parentsand the productive tilling capacity of female lines (Table 1).

**Table 1.**
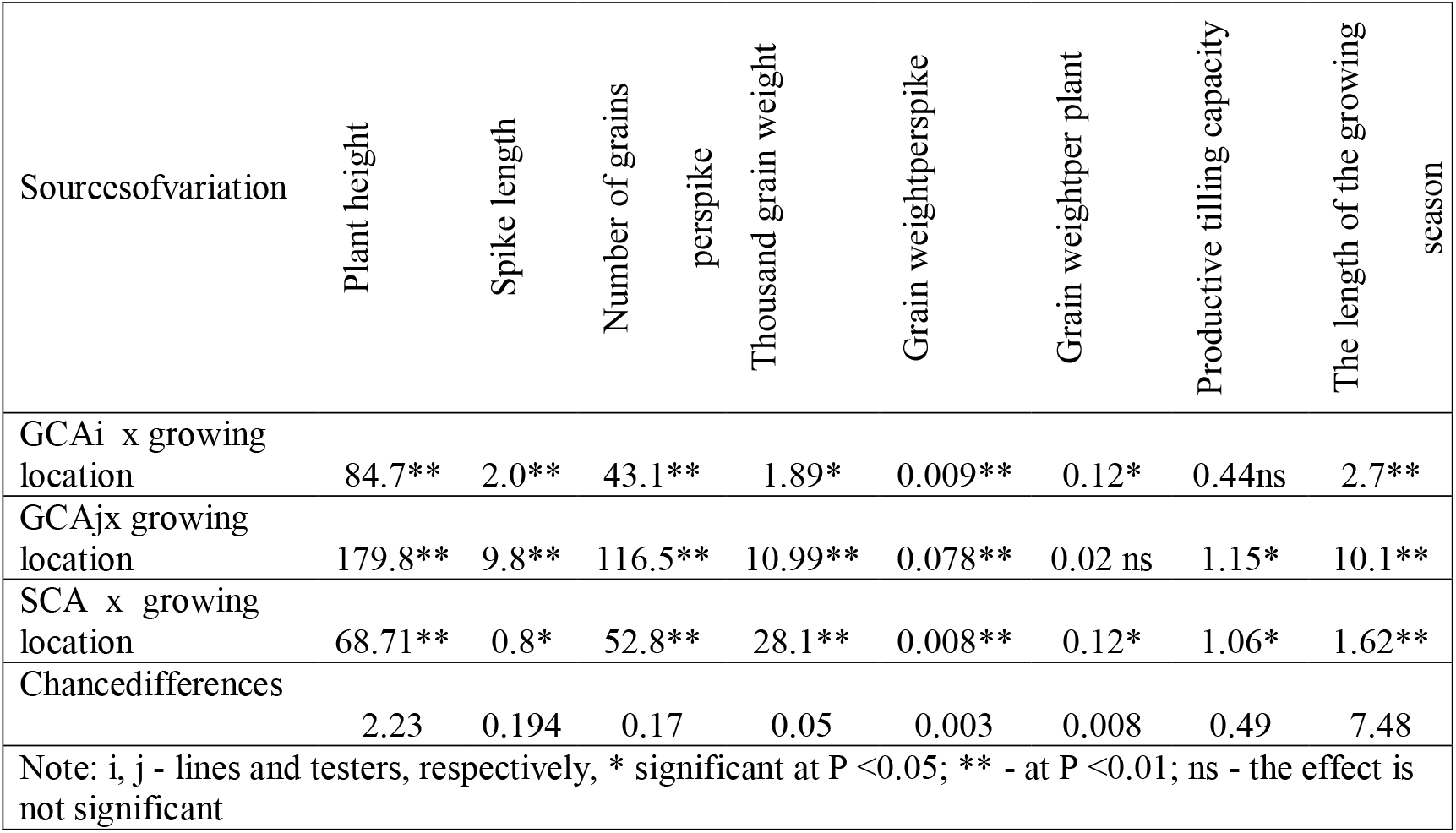
Analysis of variance of combining ability and its interaction with the growing locationof hybrids

Analysis of variance revealed significant differences in the GCA and SCA of the traits under study (Fact>Ftabl), which made it possible to proceed to the estimatesof the GCA and SCAeffects.

The determination of the studied traits involved additive (GCA) and non-additive effects (SCA), and the predominance of certain types of gene interactions depended on the cultivation conditions (Table 2).

**Table 2.** Analysis of variance of combining ability of the traits under study at two testing sites

| Sources of variation | Plant height | Spike length | Number of grains per spike | Thousand grain weight | Grain weight per spike | Grain weight per plant | Productive tillering capacity | The length of the growing season |
| --- | --- | --- | --- | --- | --- | --- | --- | --- |
| *Rice systems of the Kyzylorda region (saline soils)* |  |  |  |  |  |  |  |  |
| GCAi | 344.6** | 21.9** | 67.0** | 36.43** | 0.144** | 0.37* | 2.1 ns | 175.6* |
| GCAj | 103.2** | 14.8** | 178.0** | 51.19* | 0.245** | 0.35* | 2.4* | 712.5** |
| SCA | 110.2** | 5.9* | 37.3* | 696.6** | 0.132** | 0.21 ns | 2.1* | 725.8** |
| Proportion of GCA % | 80 | 87 | 87 | 11 | 65 | 55 | 65 | 77 |
| *Piedmont zone of the Almaty region (nonsaline soils)* |  |  |  |  |  |  |  |  |
| GCAi | 195.9* | 14.3* | 125.3** | 58.9** | 0.152* | 0.24* | 1.8* | 170.0* |
| GCAj | 814.5** | 50.25** | 200.5** | 84.0** | 0.453** | 0.23* | 1.4 ns | 622.7** |
| SCA | 633.3** | 24.7** | 196.5** | 40.5** | 0.227** | 0.18 ns | 1.3 ns | 804.3** |
| Proportion of GCA % | 61 | 72 | 59 | 78 | 73 | 72 | 71 | 49 |
| Note: *significant at P < 0.05; ** significant at P < 0.01; ns - the effect is not significant |  |  |  |  |  |  |  |  |

Regardless of the growing location, additive effects prevailed in the determination of such traits as plant height, spike length, number of grains per spike, grain weight per spike, productive tilling capacity. And in the demonstration of the trait “thousand grain weight” in the rice system conditions the Kyzylorda region, the main influence was exerted by the nonalleliceffects of genes, in contrast to the piedmont zoneconditions of the Almaty region. If, in the conditions of the Kyzylorda region, the allelic effects of genes exceeded the nonallelic onesin the length of the growing season, in the piedmont zoneof the Almaty region this trait was equally controlled by both additive and non-additive gene interactions.

The proportion of the additive effects of the genes of seed parentsalso depends on the influence of the environment. In contrast to thepiedmont zone of the Almaty region, in the conditions of the rice systems of the Aral Sea region, the mean squares of the GCA of the female lines exceeded the GCA values of the testers, being inferior in terms of theSCA in the plant height and spike length. The number of grains per spike, thousand grain weight, grain weightperspike, and the length of the growing season in both zones were mainly determined by the additive genes of male parents; however, in the conditions of rice systems, its slight decrease was observed. The cultivation zones had a significant impact on thedemonstrationof the specific combining ability of the thousand grain weightand the length of the growing season. In the conditions of saline soils of the Aral Sea region, the thousand grain weight, in thepiedmontzone of the Almaty region, the length of the growing season was noted by more pronounced nonallelic interactions.

### Plant height

A distinctive feature of the Syrian barley samples is their dwarfness (not higher than 50 cm), as the most possible genetic sources of short-stature, which is of certain interest for breeding varieties in the conditions of the irrigated zone of the Almaty region. The best in breeding terms are the varieties WI2291/Roho/WI2269and Rihane, distinguished by persistent low GCA in height and positive in the number of grains, thousand grain weight,grain weightperspike, in combination with low GCA effects in the length of the growing season simultaneously in two zones (sources of early maturity). Variety-testers Saule, Donetskiy 8, have a fairly low GCA in height, but in the course of further analysis of the combining ability, their negative impact on all traits of productivity have been revealed. However, as studies by Al-Imran Dianga [14] have shown, the results of one population cannot be extrapolated to another population. The effect of the gene must be assessed for each population, that is, the values of the specific combining ability must be assessed and taken into account.

One of the determining factors in the zoning of a particular variety of barley in the conditions of the rice crop rotation of the Kyzylorda region is the height of the plants, since it is mainly cultivated as a cover crop of perennial grasses. Therefore, the development of varieties with optimal stem sizes (not lower than 75 cm), combining early maturity to avoid overgrowth of grasses above, is an urgent direction in barley breeding for this region.

Considering the greatest practical value of these traits, the best seed parentsshould be recognized as the Odessa 100 and Harmal varieties, in which a very favorable combination of high GCA and SCAhas been noted, besides the additive variance in quantitative terms somewhat has predominated over the non-additive one, which indicates the possibility of positive transgressions in subsequent generations.

### Spike length

Novarieties with stable highestimates of the GCAwere identified simultaneously at two sites according to the spike length. We can note Harmal, which is characterized by relatively stable positive estimates, that is, independent of the environmental differences of high and average GCA. In the conditions of the Aral Sea region, the Odessa 100 variety, which also has high variances of the SCA, can be used in heterosisand linear selection to improve this trait (Table 4).

**Table 4.** Estimates of the GCA effects and variance of the SCA of varieties by morpho-biological traits, depending on the testing sites

| Lines, varieties | Site | Plant height |  | Spike length |  | The length of the growing season |  |
| --- | --- | --- | --- | --- | --- | --- | --- |
|  |  | GCA | SCA | GCA | SCA | GCA | SCA |
| WI2291/Roho/WI2269 | A | - 8.6 | 30.93 | - 2.5 | 4.9 | - 2 | 95.2 |
|  | B | - 12.4 | 280.5 | + 0.3 | - 0.9 | - 4 | 125.2 |
| WI2291*2/WI2269 | A | + 0.2 | - 4.4 | + 1.3 | 7.1 | + 1 | 71.3 |
|  | B | + 4.4 | 123.2 | - 2.6 | 0.5 | + 3 | 101.3 |
| Rihane | A | - 6.6 | 160.9 | - 1.8 | 4.7 | - 10.5 | 160.5 |
|  | B | - 1.4 | 270.1 | + 1 | 16.9 | - 7.5 | 190.5 |
| Harmal | A | + 15 | 210.5 | + 3 | 2.8 | + 3.5 | 125.0 |
|  | B | + 9.4 | 164.5 | + 1.3 | 2.5 | + 8.5 | 155.5 |
| HCP<sub>05</sub> |  | 7.8 | 146.7 | 2.9 | 8.7 | 5.0 | 63.3 |
| Saule | A | - 3.4 | 23.3 | - 1 | 3.8 | - 3 | 135.7 |
|  | B | + 1.6 | 46.7 | - 1 | 2.6 | - 3 | 165.5 |
| Donetskiy8 | A | - 2.4 | 50.2 | - 1.2 | 2.8 | + 10.4 | 71.0 |
|  | B | - 4 | 141.9 | + 2 | 2.7 | + 0.6 | 101.6 |
| Odessa 100 | A | + 5.8 | 136.6 | + 2.2 | 6.2 | + 0.6 | 72.4 |
|  | B | + 2.4 | 142.0 | - 1 | 5.8 | + 2.4 | 102.3 |
| HCP<sub>05</sub> |  | 2.1 | 82 | 0.9 | 2.9 | 7.6 | 73.6 |
| Note: Sites: A – Kyzylorda region; B – Almaty region |  |  |  |  |  |  |  |

### Number of grains perspike

In the conditions of the rice systems of the Aral Sea region, the lines WI2291*2/WI2269 andHarmal, and tester Odessa 100 were identified according to GCA, which were also characterized by high valuesin the “number of grains per spike” (on average 22–24 pcs.) in comparison with other genotypes being studied. The lines Rihane and Harmal and testers Odessa 100 andSaule are distinguished by highvariances of the SCA, which are recommended to be used to obtain certainheterotic combinations.

In specific combinations, the best were the hybrids WI2291*2/WI2269 x Odessa 100; WI2291/Roho/WI2269 x Saule; WI2291*2/WI2269 x Donetskiy 8; Harmal x Saule. In the conditions of the piedmont zoneof the Almaty region, positive estimates of the GCA effects were noted for the line WI2291/Roho/WI2269 and Rihane. According to this trait, there is no presence of varieties stable in terms of GCA; only Odessa 100 can be identified, distinguished by positive high and average estimates of the GCA effects, which is also characterized by high values in the “number of grains per spike” (on average 22-24 pcs.) in comparison with other genotypes being studied, that is, the greatest expression of the trait dominates, therefore, it has the largest number of alleles that positively determine the trait (Table 5).

**Table 5.**
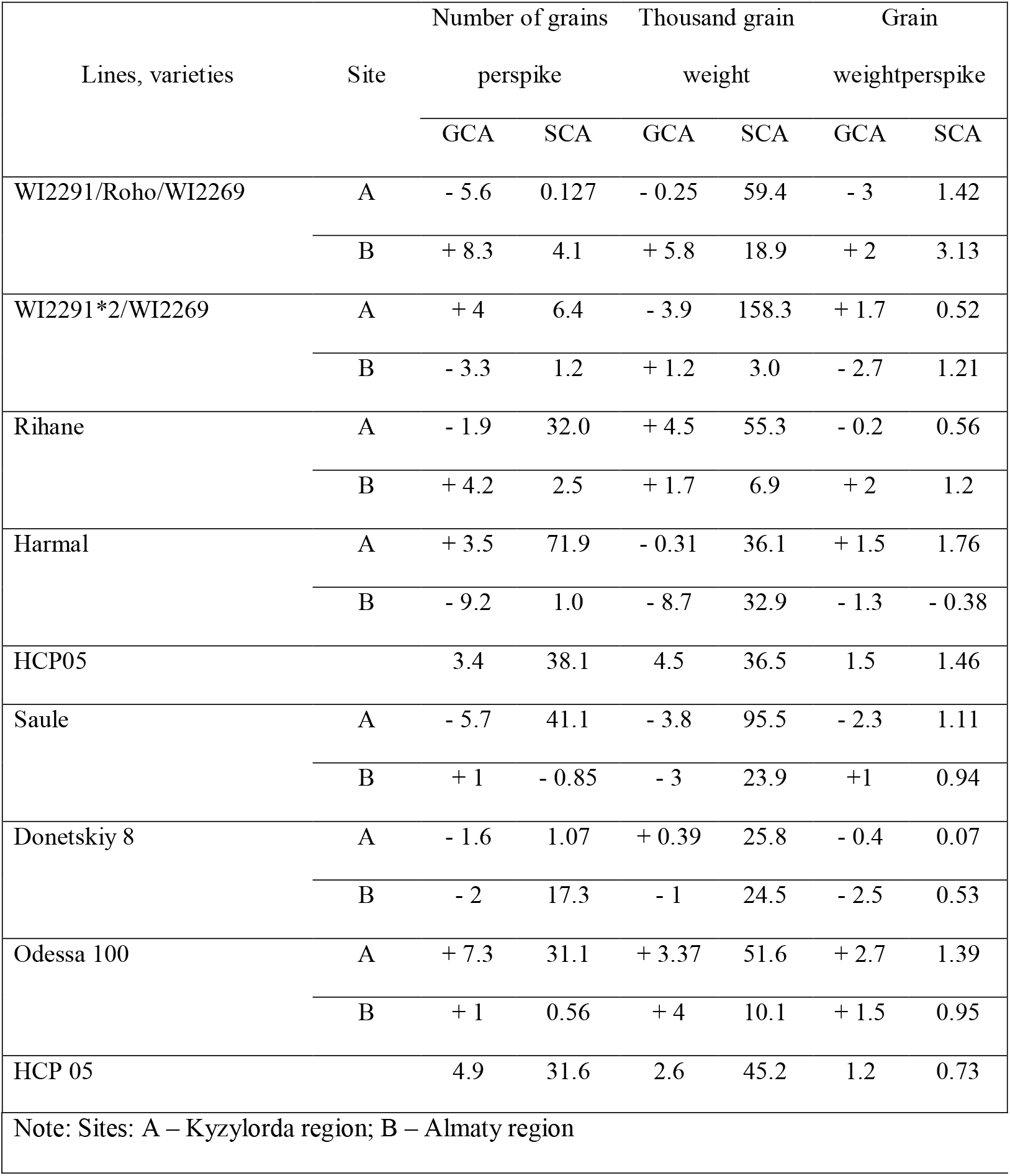
Estimates of the GCA effects and variance of the SCA of varieties by traitsof productivity, depending on the testing sites

| Lines, varieties | Site | Number of grains |  | Thousand grain |  | Grain |  |
| --- | --- | --- | --- | --- | --- | --- | --- |
|  |  | perspike |  | weight |  | weightperspike |  |
|  |  | GCA | SCA | GCA | SCA | GCA | SCA |
| WI2291/Roho/WI2269 | A | - 5.6 | 0.127 | - 0.25 | 59.4 | - 3 | 1.42 |
|  | B | + 8.3 | 4.1 | + 5.8 | 18.9 | + 2 | 3.13 |
| WI2291*2/WI2269 | A | + 4 | 6.4 | - 3.9 | 158.3 | + 1.7 | 0.52 |
|  | B | - 3.3 | 1.2 | + 1.2 | 3.0 | - 2.7 | 1.21 |
| Rihane | A | - 1.9 | 32.0 | + 4.5 | 55.3 | - 0.2 | 0.56 |
|  | B | + 4.2 | 2.5 | + 1.7 | 6.9 | + 2 | 1.2 |
| Harmal | A | + 3.5 | 71.9 | - 0.31 | 36.1 | + 1.5 | 1.76 |
|  | B | - 9.2 | 1.0 | - 8.7 | 32.9 | - 1.3 | - 0.38 |
| HCP05 |  | 3.4 | 38.1 | 4.5 | 36.5 | 1.5 | 1.46 |
| Saule | A | - 5.7 | 41.1 | - 3.8 | 95.5 | - 2.3 | 1.11 |
|  | B | + 1 | - 0.85 | - 3 | 23.9 | +1 | 0.94 |
| Donetskiy 8 | A | - 1.6 | 1.07 | + 0.39 | 25.8 | - 0.4 | 0.07 |
|  | B | - 2 | 17.3 | - 1 | 24.5 | - 2.5 | 0.53 |
| Odessa 100 | A | + 7.3 | 31.1 | + 3.37 | 51.6 | + 2.7 | 1.39 |
|  | B | + 1 | 0.56 | + 4 | 10.1 | + 1.5 | 0.95 |
| HCP 05 |  | 4.9 | 31.6 | 2.6 | 45.2 | 1.2 | 0.73 |

|  |
| --- |
| Note: Sites: A – Kyzylorda region; B – Almaty region |

### Thousand grain weight

In the conditions of the Kyzylorda region, the lines Rihane andthe Odessa 100 varietywere identified according to the GCA; WI2291*2/WI2269 – according to the SCA, which can be used to highlight certain successful combinations when crossing by this trait.

The piedmont zone of the Almaty region determined the high combination value of the lines WI2291/Roho/WI2269, in contrast to the Aral Sea zone, which negatively affected the grain size. Regardless of the growing areas, estimates of the GCA effectsin the Odessa 100 variety are highly significant, in combination with a high SCA in the Aral Sea zone, which is of great value for both synthetic and heterosisselection.In specific combinations, the best were the hybrids: WI2291*2/WI2269XDonetskiy 8; WI2291*2/WI2269 xOdessa 100; Harmal X Saule.

### Grain weightperspike

The conditions of the saline soils of the Aral Sea region had a negative effect on the GCA of varieties WI2291/Roho/WI2269, Saule, Donetskiy 8, Saule in terms of grain weight per spike, but did not change the estimates of the Odessa 100 variety. Whereas, the conditions of the piedmont zone, on the contrary, had a positive effect on the GCA of the lines WI2291/Roho/WI2269 andRihane, which also had high GCA effects in thousand grain weightand number of grains per spike, respectively. The hybrids obtained on their basis are characterized by high productivity. Special attention should be paid to the samples Harmal and WI2291/Roho/WI2269, which have been distinguished by a high variance of the SCAin the rest traits of productivity, which can be recommended for obtaining promising combinations for the selection of transgressive lines.

Thus, varieties and lines stable according to the GCA were determined simultaneously in two geographical points:

- Odessa 100 in two traits: the length of the growing season and thousand grain weight;
- Donetskiy 8 in two traits:the number of grains per spike and thousand grain weight;
- Saule in three traits: spike length, the length of the growing season andthousand grain weight;
- Rihane in two traits: plant height and grain weight perspike;
- WI2291/Roho/WI2269, WI2291*2/WI2269 andHarmalin three traits: plant height, spikelength and the length of the growing season.

In general, the varieties Rihane, WI2291/Roho/WI2269 and the variety-tester Odessa 100 with high GCA and SCA effects, little dependent on growing conditions, which can serve as donors of important selection parameters, are of the greatest practical interest.

## Conclusion

Genetic analysis of economically valuable traits of spring barley varieties of various ecological and geographical origin allows us to make the following conclusion:

- determination of the GCA and SCA of barley genotypes in two regions of Kazakhstan contrasting in soil and climatic conditions has revealed both the presence of a significant variation in the studied traits from the growing zones, and differences between varieties in combining ability; the role of additive and non-additive genes in the determination of the traits understudyhas been revealed;
- the role of additive genes in the genetic control of all analyzed traits is essential, therefore, the efficiency of selection and the appearance of transgressive forms in early segregating populations can be predicted for them, with the exception of the trait “thousand grain weight” and “the length of the growing season”;
- in the piedmont zone of the Almaty region, positive selection results are possible in the case of the action of both additive and non-additive effects of genes in the inheritance of plant height and the duration of the growing season, as a result of the appearance of homozygous forms;
- the salinizationconditions had a significant effect on phenotypic differences in thousand grain weight, and the piedmont zone in the length of the growing season, which resulted in an increase in the non-additive variance (SCA), i.e. in the determination of these traits, dominant and epistatic genesprevail.
- theGCA effects, in contrast to variance of the SCA, were higher and more stable in most of the traits. The changeability of variance of the SCAdepending on the years and testing sites is associated with the determination of this parameter by genes with dominant and epistatic effects, which are characterized by high sensitivity to numerous environmental factors.

In conclusion, we would like to note that the results of this research cannot fully reveal the genetic traits of objects, but they give a general idea of the inheritance of important economic and biological traits in various environmental conditions and allow breeders to build a model for future varieties and a strategy for selection work.

## Data Availability

Data used in preparing this manuscript is available from the corresponding author on reasonable request.

## Acknowledgments

This research is funded by the Ministry of Agriculture of the Republic of Kazakhstan.

## References

1. Tokhetova L. A., Umirzakov S. I., Nurymova R. D., Baizhanova B. K., Akhmedova G. B., “Analysis of Economic-Biological Traits of Hull-Less Barley and Creation of Source Material for Resistance to Environmental Stress Factors”, International Journal of Agronomy, vol. 2020, Article ID 8847753, 10 pages, 2020https://doi.org/10.1155/2020/8847753

2. Tokhetova L.A., Tautenov I.A., Zelinski G.L. (2017) Demesinova A.A. Variability of main quantitative traits of the spring barley in different environmental conditions. Ecology, Environment and Conservation, Vol 23, Issue 2, Page 1093–1098 http://www.envirobiotechjournals.com/article_abstract.php?aid=7887&iid=230&jid=3

3. NeseSreenivasulu, Andreas Graner, Ulrich Wobus, “Barley Genomics: An Overview”, International Journal of Plant Genomics, vol. 2008, Article ID 486258, 13 pages, 2008. https://doi.org/10.1155/2008/486258

4. J. A. Olfati, H. Samizadeh, B. Rabiei, Gh. Peyvast, “Griffing’s Methods Comparison for General and Specific Combining Ability in Cucumber”, The Scientific World Journal, vol. 2012, Article ID 524873, 4 pages, 2012. https://doi.org/10.1100/2012/524873

5. B. Griffing, “Concept of general and specific combining ability in relation to diallel crossing systems,” Australian Journal of Biological Sciences, vol. 9, no. 4, pp. 463–493, 1956.

6. Shawn Kaeppler, “Heterosis: Many Genes, Many Mechanisms—End the Search for an Undiscovered Unifying Theory”, International Scholarly Research Notices, vol. 2012, ArticleID 682824, 12 pages, 2012. https://doi.org/10.5402/2012/682824

7. GhislainKanfany, AmadouFofana, PangirayiTongoona, AgyemangDanquah, Samuel Offei, Eric Danquah, NdiagaCisse, “Estimates of Combining Ability and Heterosis for Yield and Its Related Traits in Pearl Millet Inbred Lines under Downy Mildew Prevalent Areas of Senegal”, International Journal of Agronomy, vol. 2018, Article ID 3439090, 12 pages, 2018. https://doi.org/10.1155/2018/3439090

8. FawziaBouchetat, AbdelkaderAissat “Evaluation of the genetic determinism of an F1 generation of barley resulting from a complete diallel cross between autochthones and introduced cultivars”, Heliyon, Volume 5, Issue 11, 2019, https://doi.org/10.1016/j.heliyon.2019.e02744

9. Emmanuel Yaw Owusu, Haruna Mohammed, KulaiAmaduManigben, Joseph Adjebeng- Danquah, Francis Kusi, Benjamin Karikari, Emmanuel Kofi Sie, “Diallel Analysis and Heritability of Grain Yield, Yield Components, and Maturity Traits in Cowpea (Vignaunguiculata (L.) Walp.)”, The Scientific World Journal, vol. 2020, Article ID 9390287, 9 pages, 2020.

10. Savchenko V.K. Genetic and statistical parameters and their use in plant breeding for productivity. - Tallinn, 1981. - P. 86–101

11. P. R. Patil, V. H. Surve, and H. D. Mehta, “Line × tester analysis in rice (Oryza sativa L.)”, Madras Agriculture Journal, vol. 99, no. 4–6, pp. 210–213, 2012. https://doc-0g-7g-docsviewer.googleus…

12. Mohamed Labdi, SamiaGhomari, SamiaHamdi, “Combining Ability and Gene Action Estimates of Eight Parent Diallel Crosses of Chickpea for Ascochyta Blight”, Advances inAgriculture, vol. 2015, ArticleID 832597, 7 pages, 2015. https://doi.org/10.1155/2015/832 597

13. M. J. Hasan, M. U. Kulsum, M. Hossain, M. H. Manzur, M. R. Mustafizur, and R. N. M. Farhat, “Combining ability analysis for identifying elite parents for heterotic rice hybrids”, Academia Journal of Agricultural Research, vol. 3, no. 5, pp. 70–75, 2015.

14. Al-Imran Dianga, Kamau W. Joseph, Ruth N. Musila, “Analysis of Combining Ability for Early Maturity and Yield in Rice (Genus: Oryza) at the Kenyan Coast”, International Journal of Agronomy, vol. 2020, Article ID 6230784, 7 pages, 2020. https://doi.org/10.1155/2020/6230784

15. Tarutina L. A., Khotyleva L. V. Assessment the variability of combinational ability in various environmental conditions / Genetic analysis of quantitative and qualitative characteristics using mathematical and statistical methods: M. Russian Academy of Agricultural Sciences All-Russian Research Institute of Technical and Economic Research, 1973-pp. 74–82

